# High-resolution single-molecule replication profiling of the human genome

**DOI:** 10.64898/2026.07.15.738623

**Authors:** Alan Tourancheau, Victoria Rojat, Diletta Ciardo, Florence Proux, Laurent Lacroix, Jean-Michel Arbona, Benjamin Audit, Olivier Hyrien, Benoît Le Tallec

**Affiliations:** IBENS, Département de biologie, École Normale Supérieure, Université PSL, CNRS, INSERM, 75005 Paris, France; IBDM, UMR7288, Case 907, Parc Scientifique de Luminy, 13288 Marseille, cedex 09, France; CNRS, ENS de Lyon, LPENSL, UMR5672, 69342 Lyon, cedex 07, France

## Abstract

Despite the considerable progress made in recent years thanks to the rise of long-read sequencing, high-resolution, single molecule (SM)-based replication profiling of large genomes has remained out of reach. Here, we present a replication labelling strategy consisting in repeated pulse-labelling of asynchronously growing cells with the thymidine analogue bromodeoxyuridine (BrdU) that greatly enhances the number of detectable replication tracks. We show that the ‘multipulse’ BrdU labelling protocol, in association with ForkML, a machine-learning method translating BrdU signals in nanopore reads into oriented and positioned replication forks, enables the establishment of a SM replication map of the entire human genome at kilobase resolution.

## Main

Single-molecule (SM) analyses of DNA replication have provided key insights into eukaryotic genome duplication, whether by uncovering fundamental principles of the replication process or by measuring crucial parameters such as fork speed or distances between origins. SM methods typically rely on the immunofluorescent detection on stretched DNA fibres of thymidine analogues incorporated into replicating DNA, which allows the visualisation of elongating forks as well as of initiation and termination events along individual molecules^1^. Albeit ideal for capturing replication flexibility and revealing intercellular heterogeneity, conventional SM techniques are unable to map replication tracks on a large scale, restricting most investigations to ‘anonymous’ forks and preventing the integration of genomic and chromatin contexts. This barrier is currently being lifted thanks in particular to the introduction of long-read sequencing-based replication mapping approaches like DNAscent^2^, FORK-seq^3^, and their derivatives^4–9^, which have replaced the immunochemical detection of thymidine analogues with direct identification by dedicated basecallers, while providing replication signal genomic coordinates. Initially established in *Saccharomyces cerevisiae* model organism, where they successfully produced whole-genome SM replication maps^2, 3^, these techniques have now been extended to higher eukaryotes^7–12^. However, the large size of metazoan genomes has thus far restrained high-coverage SM studies either to specific, enriched genomic loci^7^, or to extrachromosomal DNA^11^, with genome-wide investigations remaining cost prohibitive. SM replication profiling of the ~144 Mb Drosophila genome has lately been reported^12^, which is an important step forward although the low coverage and resolution (<10 forks per 100 kb) prevent detailed analyses. Optical mapping of DNA replication^13–16^ constitutes a valuable alternative for high-throughput SM studies of metazoan genomes thanks to ultra-long, mappable DNA molecules, as illustrated by the genome-wide identification of early-firing initiation sites in human cells^16^. Limitations include an applicability essentially confined to cell extracts and synchronised cells due to the use of fluorescent nucleotides that do not passively enter cells, and a maximal resolution of ~15 kb for initiation site location.

In order to map replication fork progression in the human genome, we have recently introduced ForkML, a machine learning method interpreting signals of the thymidine analogue bromodeoxyuridine (BrdU) in nanopore reads of genomic DNA from asynchronous cells exposed to two successive BrdU pulses separated by cell incubation in BrdU-free medium^9^. This procedure captures an ongoing replication fork position at two timepoints, providing access to fork speed by dividing the genomic distance travelled by the fork between pulses by the duration of the inter-pulse time interval. Since our labelling protocol does not use a thymidine chase step that would interfere with subsequent BrdU incorporation, we reasoned that, in principle, an unlimited number of consecutive BrdU pulses could be applied. Here, we show that multiplying BrdU pulses enables fork tracking throughout S phase and proportionally increases the number of detectable replication signals, thereby alleviating coverage limitations for large genomes. Combined with ForkML, this ‘multipulse’ strategy allows the computation of a high-resolution SM replication profile of the entire human genome.

We aimed to apply the ‘multipulse’ approach to the commonly used HCT116 epithelial cell line. BrdU pulse duration and concentration were set to 4 min and 10 μM, respectively, with a 30-min interval between pulse starts, as previously defined^9^. We decided to perform 24 consecutive BrdU pulses so that the total duration of the experiment (i.e. 12 h) would match HCT116 median doubling time (Supplementary Fig. 1a, b). Therefore, all cells would inevitably pass through S phase during this period and duplicate their genome in the presence of BrdU. Specifically, the labelling protocol consisted in pulsing exponentially growing cells with BrdU, washing them twice with prewarmed PBS, incubating them in BrdU-free conditioned medium until 30 min after the pulse start, then repeating this operation 23 times before harvesting the cells (Fig. 1a). Conditioned medium was taken from a ‘sister’ culture dish containing an equivalent number of cells and plated at the same time as the labelled one; it was used instead of fresh medium to ensure that the medium composition remained consistent in the course of the experiment. Importantly, we verified that cell incubation with 10 µM BrdU had no detectable effect on HCT116 cell growth for up to 48 h (Supplementary Fig. 1c). Flow cytometry analysis further showed that the ‘multipulse’ procedure had only a marginal impact on cell distribution in the cell cycle, with multi-labelled cells merely exhibiting a slight depletion in G1 and accumulation in G2/M (Supplementary Fig. 1d, e). Moreover, this likely resulted from the repeated washes and medium switches rather than from BrdU labelling itself, as indicated by the comparable cell cycle distribution of cells subjected to 24 pulses with PBS, the BrdU solvent, alone (Supplementary Fig. 1d, e). The repeated exposure of cells to BrdU did not lead to phosphorylation of the checkpoint protein Chk1 (Supplementary Fig. 1f), suggesting that no replication stress was being induced within the timeframe of the experiment. The main drawback of our labelling protocol was a ~50% cell loss due to the frequent changes in culture medium (Supplementary Fig. 1g, h). Altogether, these results show that the ‘multipulse’ strategy allows the examination of replication dynamics under physiologically relevant conditions.

**Figure 1.**
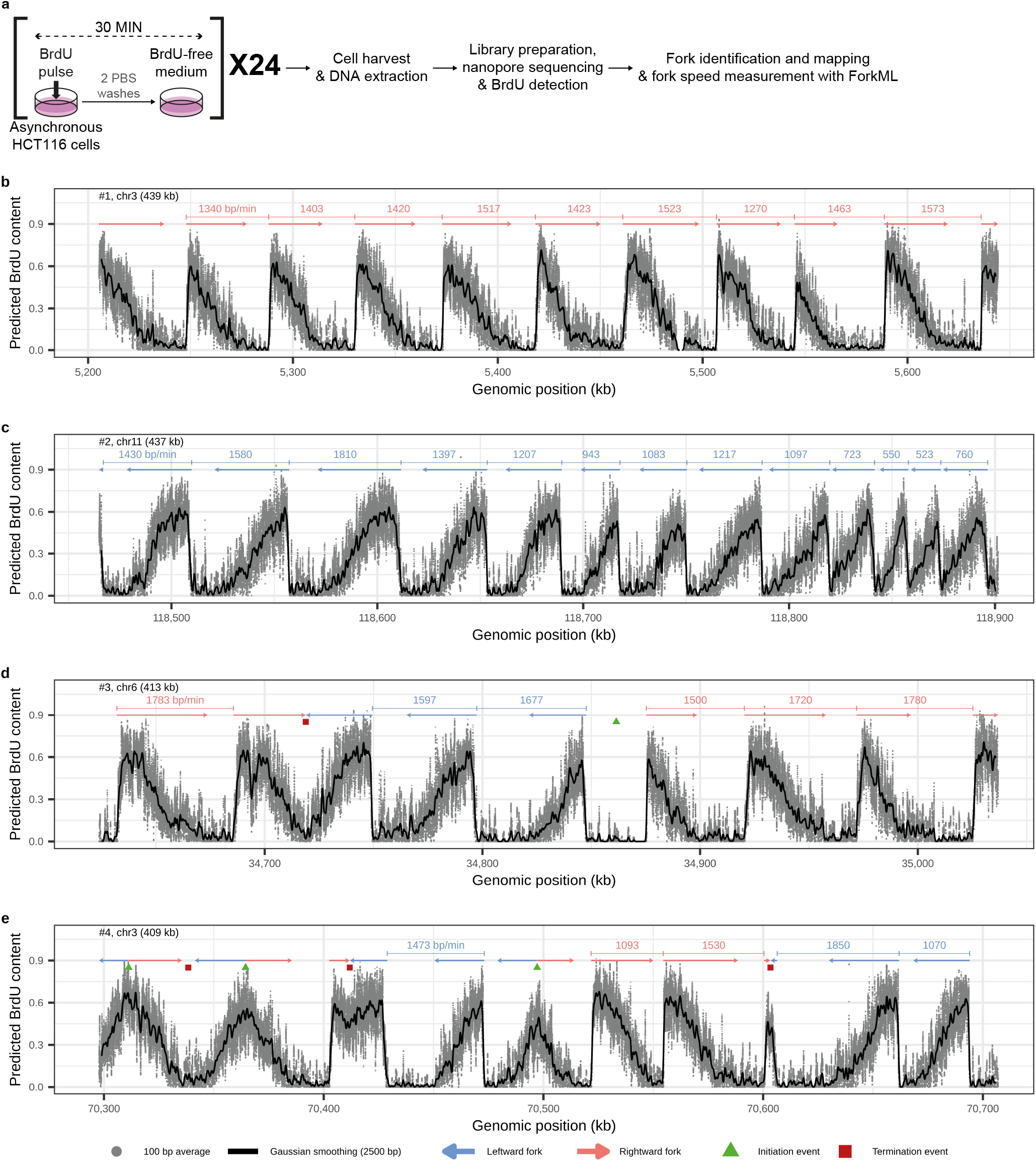
Analysis of the replication dynamics of the human genome using ‘multipulse’ BrdU labelling and ForkML. **a**, Experimental workflow. BrdU pulse concentration and duration were set to 4 min and 10 μM, respectively. **b-e**, Examples of BrdU content profiles of nanopore reads of genomic DNA from HCT116 cells pulse-labelled 24 times with BrdU. ForkML-inferred replication fork footprints are shown above each molecule, with blue and red arrows indicating leftward and rightward forks, respectively. Green triangles and red squares mark ForkML-inferred positions of initiation (diverging forks) and termination (converging forks) events, respectively. Fork speed estimates by ForkML are indicated on top, where available. Grey points, BrdU signal averaged over 100 bp; black line, Gaussian-smoothed signal over 2.5 kb.

‘Multipulse’ replication tracks appeared mainly as series of adjacent, similar, asymmetric BrdU signals (Fig. 1b-e). Each consisted of a steep upward segment starting from the baseline followed by a gradual downward section converging towards background BrdU levels, which, as demonstrated earlier^9^, corresponded to BrdU incorporation during the pulse and during cell incubation in BrdU-free medium, respectively, and revealed fork orientation. In other words, one asymmetrical BrdU signal represented an elongating fork progressing in a given direction over a period of 30 min. For example, in Fig. 1b, read#1 shows a rightward-moving fork captured eleven times at 30-min intervals, travelling over 400 kb in the meantime; read#2 (Fig. 1c) displays a leftward fork seized fourteen times in a row, that is, throughout the entire S phase, estimated at 6.6 h in HCT116 cells (Supplementary Fig. 1b, e). Initiation events were identified both in the form of diverging forks (Fig. 1d) and of symmetrical BrdU signals corresponding to pairs of forks initiated after the start of the pulse (Fig. 1e); likewise, termination events were detected either as separate, converging forks (Fig. 1d) or as symmetrical BrdU signals representing fork merging before the return of the BrdU level to the baseline (Fig. 1e). Remarkably, the ‘multipulse’ approach made it possible to track a fork from the moment it emanated from an origin to the time it met a converging fork (Fig. 1d, e), thereby showing the DNA replication programme unfold before our very eyes.

Using the ForkML pipeline^9^, we identified ~12 times more oriented replication tracks in HCT116 cells pulsed 24 times than in cells pulsed only twice (10,172 versus 818 tracks per Gb of mapped DNA on average; see Methods), representing an increase proportional to that in the number of BrdU pulses, as anticipated. Eleven ‘multipulse’ experiments sequenced using Oxford Nanopore Technologies (ONT) PromethION device each yielded between 300,000 and one million oriented replication tracks, depending on the sequencing throughput (Table 1), which allowed the computing of a SM-based replication fork directionality (RFD) profile of the human genome at 1 kb resolution (Fig. 2a and Supplementary Fig. 2). This profile showed the distinctive alternation of regions with predominant rightward and leftward fork progression, as recurrently observed^17–20^, with left-to-right and right-to-left transitions indicating initiation and termination zones, respectively. Above all, it was strikingly similar to that generated in the same cell line using the GLOE-seq technique^18^ (Fig. 2b, c; Spearman’s pairwise correlation coefficient of 0.82), which validated the robustness of our approach. The main difference lay in the higher amplitude of the ‘multipulse’ RFD profile, often reaching absolute values close to 1 indicating unidirectional replication across the whole cell population, whereas GLOE-seq absolute RFD values hardly exceeded 0.7; this is likely attributable to a higher background noise in the GLOE-seq profile.

**Table 1.** Detailed information about the ‘multipulse’ samples sequenced in this study. Rep, replicate.

| Run name | Mapped DNA (Gb) | Number of oriented replication tracks | Number of initiation events | Number of termination events |
| --- | --- | --- | --- | --- |
| HCT116_rep1 | 31.7 | 333,303 | 47,289 | 47,677 |
| HCT116_rep2 | 29.2 | 310,858 | 43,508 | 43,483 |
| HCT116_rep3 | 91.3 | 974,222 | 126,849 | 126,695 |
| HCT116_rep4 | 73.4 | 728,850 | 102,855 | 100,715 |
| HCT116_rep5 | 97.1 | 1,016,387 | 146,187 | 146,056 |
| HCT116_rep6 | 73.6 | 762,719 | 112,154 | 111,890 |
| HCT116_rep7 | 103.5 | 1,019,308 | 148,298 | 145,928 |
| HCT116_rep8 | 100.6 | 1,022,747 | 148,359 | 147,541 |
| HCT116_rep9 | 81.0 | 774,826 | 110,925 | 108,706 |
| HCT116_rep10 | 90.9 | 912,267 | 131,766 | 129,738 |
| HCT116_rep11 | 107.9 | 1,044,768 | 139,865 | 137,481 |
| *Total* | *880.2* | *8,900,255* | *1,258,055* | *1,245,910* |

**Figure 2.**
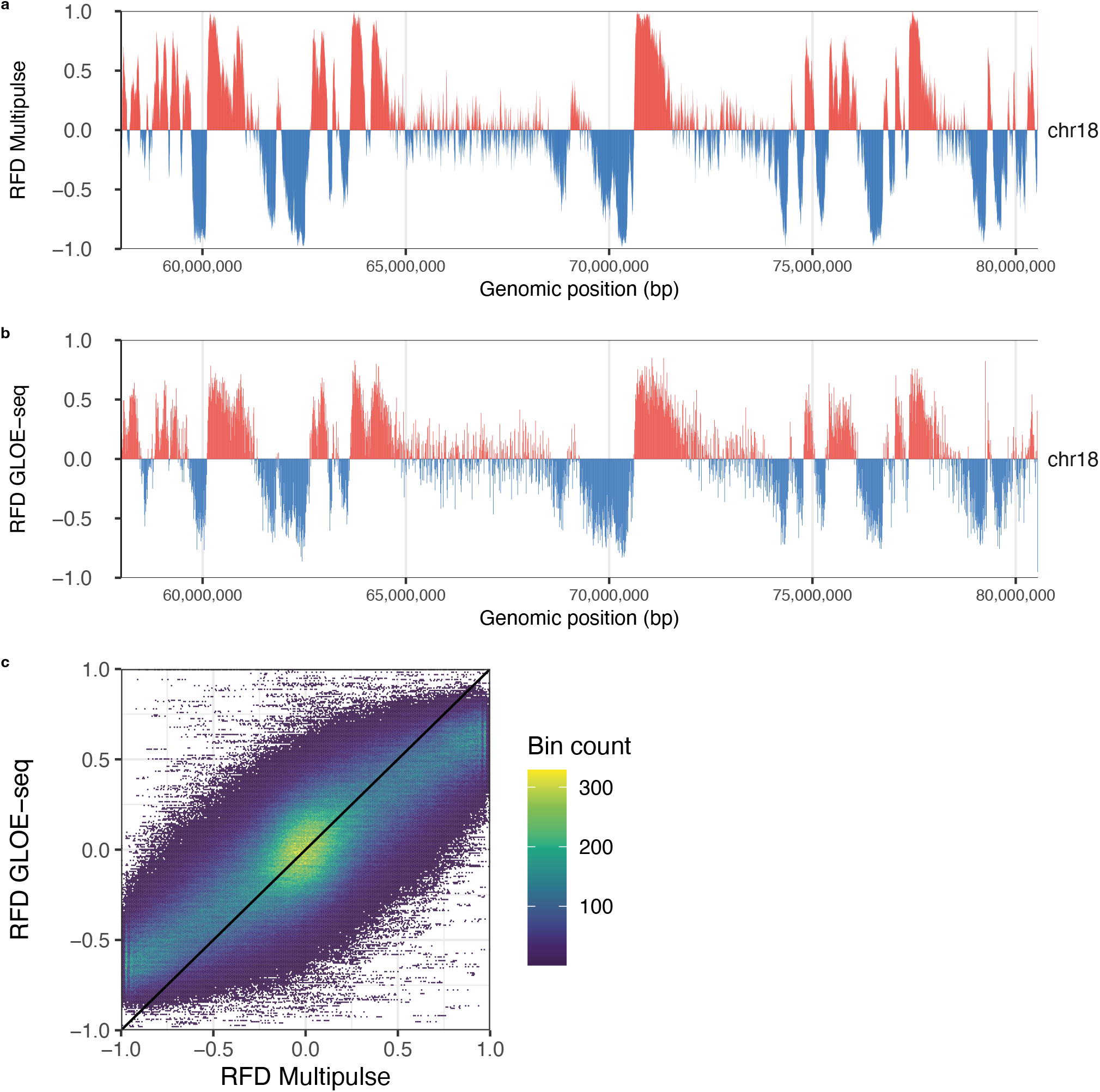
High-resolution, SM-based RFD profiling of the human genome. **a**, RFD profile in 1 kb windows of a ~20 Mb section of chromosome 18 built upon aggregation of oriented replication tracks detected by ForkML in nanopore reads of genomic DNA from HCT116 cells pulse-labelled 24 times with BrdU. Mean coverage of ~65x. All human chromosomes are shown in Supplementary Fig. 2. **b**, GLOE-seq RFD profile in HCT116 cells of the same chromosomic region as in **a** computed in 10 kb windows. Mean coverage of ~1153x. **a, b**, Positive (predominance of rightward forks) and negative (predominance of leftward forks) RFD values are in red and blue, respectively. **c**, Genome-wide comparison between ‘multipulse’ and GLOE-seq RFD profiles represented as a 2D density plot using hexagonal bins. The diagonal indicates equal RFD values between the two methods. GLOE-seq data in **b, c** are from ref. 18.

In addition to establishing a high-resolution SM replication map of the human genome, the ‘multipulse’ strategy identified over one million individual initiation and termination events (Table 1), which is more than two orders of magnitude higher than recent SM studies of metazoan genomes^7, 8, 11, 12^. It also rivals the ~1 million early-initiation events positioned using optical replication mapping^16^, while locating origin firing throughout S phase. Finally, like the ‘double-pulse’ labelling procedure^9^, the ‘multipulse’ protocol makes it possible to monitor fork velocity, and can do so over much longer periods – routinely several hours, or even the entire S phase duration. For instance, the speed of the long-travelling rightward fork shown on read#1 of Fig. 1b could be computed over nine successive 30-min periods simply by calculating the ratio between the distance separating fork positions at the start of two given, consecutive BrdU pulses and the duration of the time interval. The fork in question exhibited a fairly uniform movement during the 4.5-hour monitoring period (range of 1.3-1.6 kb/min, median of 1.4 kb/min), in sharp contrast with the leftward fork presented on read#2 of Fig. 1c, which substantially accelerated in the course of S phase (speed measured 13 times; range of 0.5-1.8 kb/min, median of 1.1 kb/min). ‘Multipulse’ experiments are therefore particularly well-suited to an in-depth analysis of fork progression dynamics, shedding light on the consistency, or lack thereof, of fork movement during S phase.

In conclusion, our ‘multipulse’ approach combines the ability to generate high-resolution, genome-wide replication profiles of large genomes with the capacity to measure fork speed throughout S phase and precisely locate individual initiation and termination events on an unprecedented scale. It fully bridges the gap between SM- and population-based assays and finally brings SM analyses of DNA replication into the genomic era, with the power to reveal the replication strategy of the human genome.

## Methods

### Cell line and cell culture

HCT116 human colon carcinoma cells, purchased from the American Type Culture Collection (ATCC; #CCL-247), were grown in McCoy’s 5A medium (Gibco #16600082 and #26600023) with 10% fetal bovine serum (Dominique Dutscher #500105G1G, batch P190801, and #500105M1M, batch S00G0), supplemented with 100 U.ml^-1^ penicillin and 100 µg.ml^-1^ streptomycin (Sigma-Aldrich #P4333). Cells were grown at 37 °C, 20% O_2_, 5% CO_2_, and routinely confirmed to be negative for mycoplasma contamination.

### Multiple BrdU pulse-labelling of neosynthesised DNA

Exponentially growing cells were pulse-labelled for 4 min with 10 µM BrdU (Sigma-Aldrich #B5002, dissolved in PBS), washed twice with prewarmed PBS, incubated in BrdU-free conditioned medium until 30 min after the start of the BrdU pulse, and the whole process was repeated 23 times, for a total experiment duration of 12 h (24*30 min). Cells were then washed with PBS, harvested by trypsinisation, and counted before genomic DNA was extracted using Monarch HMW DNA Extraction Kit for Cells & Blood (New England Biolabs #T3050) according to the manufacturer’s instructions for ultra-long DNA sequencing applications. Conditioned medium came from a ‘sister’ culture dish containing cells plated at the same time and in equal number to those labelled; as many ‘sister’ culture dishes were prepared as there were BrdU pulses. Control dishes with untreated cells were systematically used to assess cell loss during ‘multipulse’ experiments (Supplementary Fig. 1g); the number of cells recovered after 24 pulse-labelling cycles with either BrdU or PBS was also evaluated (Supplementary Fig. 1h).

### Cell growth and doubling time

Cell number was determined at regular intervals over periods of 72 to 168 h. Doubling time (T) of exponentially growing cells was estimated on the basis of the growth curves according to the formula T = [culture duration*log(2)]/[log(final number of cells) − log(initial number of cells)].

### Cell cycle analysis

Exponentially growing, untreated cells or cells pulse-labelled 24 times with BrdU or PBS as described above were incubated with 10 μM 5-ethynyl-2’-deoxyuridine (EdU, Jena Bioscience #CLK-N001-100) for 1 h and fixed in ethanol. Fixed cells were pelleted, incubated in PBS-0.2% Triton X-100 for 20 min at room temperature, washed with PBS-0.1% BSA, and subjected to ‘Click Chemistry’ to fluorescently label EdU with AF647 dye by incubation in PBS supplemented with 20 µM AF647-Picolyl-Azide (Jena Bioscience #CLK-1300A-1), 2 mM CuSO4 (Jena Bioscience #CLK-MI004-50), and 10 mM Na-Ascorbate (Jena Bioscience #CLK-MI005-1G) for 30 min at room temperature (protected from light), before a final wash with PBS-0.1% BSA. DNA was counterstained with 4′,6-diamidino-2-phenylindole (DAPI, Millipore #5.08741.0001). Samples were analysed using a ZE5 Cell Analyzer (Bio-Rad) with Everest software (version 3.2.12.0). Data were processed using FlowJo v10.10.1.

### Immunoblotting

Total protein extracts of untreated cells, cells pulse-labelled 24 times with BrdU (as described above), or cells treated with 2 mM hydroxyurea (Sigma-Aldrich #H8627) for 24 h were obtained using RIPA buffer (150 mM NaCl, 1% NP-40, 0.1% SDS, 1% sodium deoxycholate, 50 mM Tris-HCl pH 7.5, 1 mM EDTA, and 1 mM EGTA) with Pierce Universal Nuclease for Cell Lysis (ThermoFisher Scientific #88700) plus protease and phosphatase inhibitors (PMSF 200 µM; Protease Inhibitor Cocktail, Sigma-Aldrich #P8340; Phosphatase Inhibitor Cocktail 2, Sigma-Aldrich #P5726). Briefly, cells were incubated in RIPA buffer for 20-30 min on ice, centrifugated for 10 min at 4°C at 14000 rpm, and the protein-containing supernatant was collected. Protein concentration was measured using the Bradford assay. 33-50 μg of proteins were separated by SDS-PAGE on a 12% gel and transferred to a nitrocellulose membrane. Chk1, phospho-Chk1 (Ser345), and ß-actin immunoblots were performed with a mouse anti-Chk1 antibody at 1:1000 (Santa Cruz #sc-8408, batch 1009), a rabbit anti-phospho-Chk1 (Ser345) antibody at 1:1000 (Cell Signaling Technology #2348, batch 18), and a mouse anti-ß-actin antibody at 1:1000 (Cell Signaling Technology #58169, batch 1), respectively, using HRP-conjugated anti-rabbit (Promega #W401B) or anti-mouse (Promega #W402B) at 1:50000 as secondary antibodies. Detection was performed with SuperSignal West Femto Maximum Sensitivity Substrate (ThermoFisher Scientific #34095) chemiluminescent reagents. Amersham ImageQuant 800 (GE, software version 1.1.2) was used for imaging.

### Library preparation and data acquisition

Samples were sequenced using R9.4.1 PromethION flow cells from ONT. Sequencing libraries were prepared with ONT ligation sequencing kit SQK-ULK001 according to ONT protocol. During sequencing, flow cells were washed with ONT EXP-WSH004 kit and libraries reloaded twice. Data were acquired using MinKNOW (ONT, versions 23.07.12 and 23.11.7) with default parameters.

### BrdU basecalling, read mapping and BrdU signal segmentation

BrdU basecalling and read mapping were performed as in ref. ^9^ (R9.4.1 datasets; Megalodon v2.2.9, Guppy v4.4.1, and minimap2 v2.26). BrdU signal segmentation was achieved with ForkML as in ref. ^9^. T2T-CHM13^21^ (v2.0) was used as reference genome.

### Comparison of the number of oriented replication tracks detected by ForkML after HCT116 cell labelling with 2 or 24 BrdU pulses

The average number of oriented replication tracks detected per gigabase of mapped DNA after cell labelling with 2 or 24 BrdU pulses was computed from the figures in Supplementary Data 1 of ref. ^9^ (HCT116_UT_R9_rep1-4 samples) and from the figures in Table 1 of this study (HCT116_rep1-11 samples), respectively.

### RFD profiles

HCT116 ‘multipulse’ RFD profile was calculated in non-overlapping 1 kb genomic windows as (RFD = (R − L)/(R + L)), where (R) and (L) are the rightward- and leftward-oriented replication track coverages within each window, respectively. Oriented replication tracks were inferred from co-directional fork footprints. For each track, fork direction was assigned to the detected segment and extended into adjacent unassigned regions up to the midpoint between neighbouring fork calls, or, at read ends, up to half the distance to the read boundary, thereby maximizing the interpretable portion of each read while avoiding overlap between conflicting fork direction assignments. All eleven HCT116 biological replicates were used. HCT116 GLOE-seq RFD profile was computed in non-overlapping 10 kb windows as described in ref. ^9^.

### Genome-wide comparison between ‘multipulse’ and GLOE-seq RFD profiles

The ‘multipulse’ and GLOE-seq RFD profiles were compared using a hexbin scatter plot. Points were summarized into hexagonal bins, the colour of which indicates the number of genomic windows of the cognate category. To account for the different resolutions (1 and 10 kb windows for the ‘multipulse’ and GLOE-seq RFD profiles, respectively), each 10 kb GLOE-seq RFD value was matched with the ‘multipulse’ RFD values of the corresponding ten 1 kb genomic windows. Spearman’s rank correlation coefficient between both profiles was calculated using the R^22^ cor function.

## Data availability

Nanopore sequencing data generated in this study will be made available upon publication. HCT116 GLOE-seq data from ref. ^18^ used in this study are available in NCBI’s BioProject database under accession code PRJNA554350 (GSM4305465 and GSM4305466).

## Acknowledgements

The authors thank IBENS GenomiqueENS facility for their assistance with nanopore sequencing and IBENS IT platform and BioClust computing cluster (Labex Memolife) for data management. This work was supported by grants from Fondation pour la Recherche Médicale [FRM EQU202203014910 to O.H.] and Agence Nationale de la Recherche [NanoPoRep ANR-18-CE45-0002, HUDROR ANR-19-CE12-0028, and SMAHGR ANR-23-CE12-0021 to B.A. and O.H.]. V.R. was supported by fellowships from the Ministère de l’Enseignement Supérieur et de la Recherche and Fondation pour la Recherche médicale [FRM FDT202404018224].

