## Supplementary information for "High-resolution single-molecule replication profiling of the human genome"

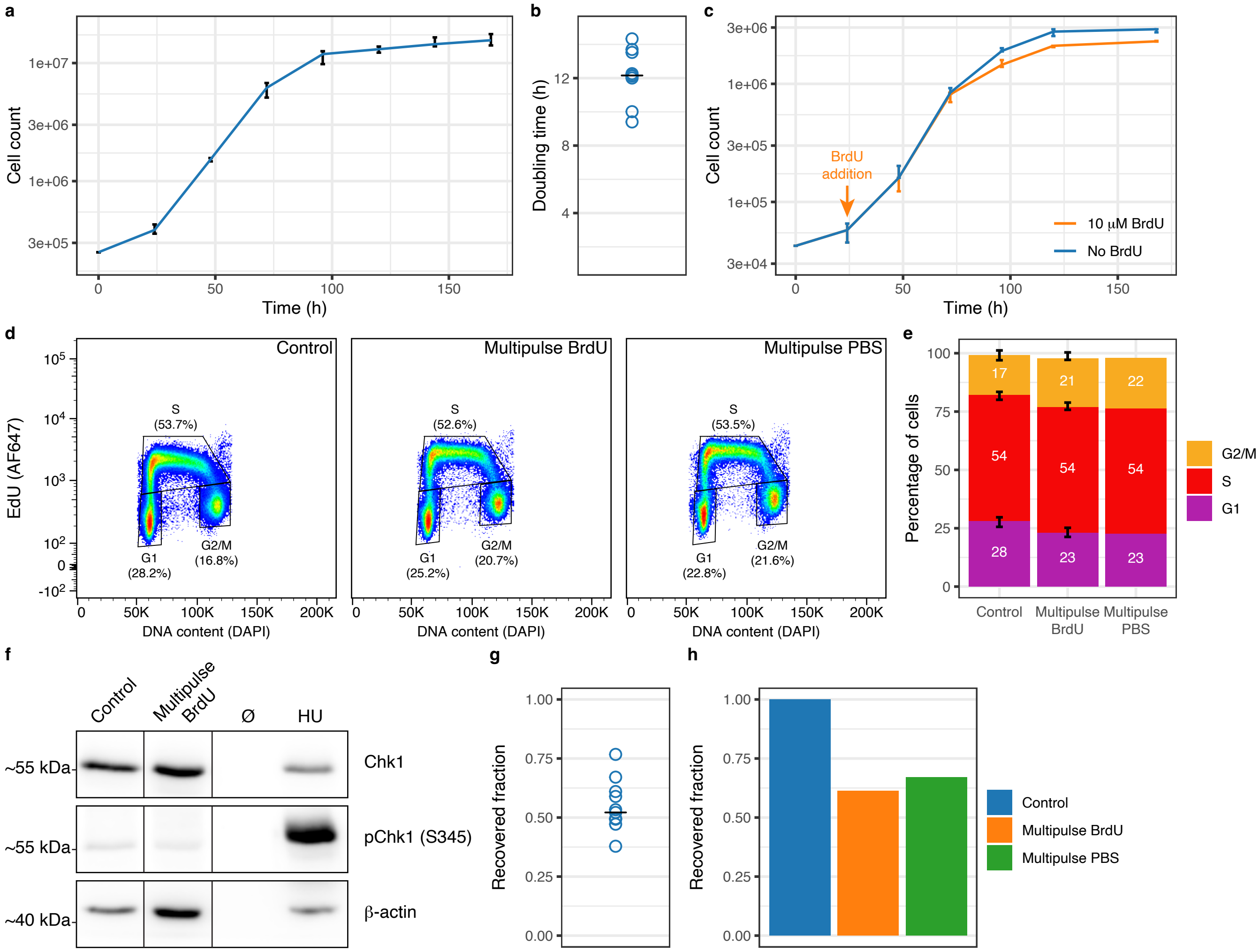


**Supplementary Figure 1. The 'multipulse' BrdU labelling strategy enables the study of replication dynamics under physiologically relevant conditions. a**, Representative growth curve of HCT116 cells. Cell counting was carried out in triplicate; median (blue curve) and extreme values (whiskers) are presented. This experiment was performed independently nine times with similar results. Cells showed exponential growth from 24 h until 72 h after plating; 'multipulse' BrdU labelling was performed after 48 h. **b**, Population doubling time of HCT116 cells. Values are population doubling times from nine independent cell cultures; HCT116 median population doubling time (black central line) is 12.2 h. **c**, HCT116 cell growth with or without 10 μM BrdU. BrdU was added 24 h after plating. Cell counting was carried out in triplicate for each condition; median (coloured curve) and extreme values (whiskers) are presented. **d**, Representative bivariate flow cytometry analysis (EdU/DNA) of the cell cycle of HCT116 cells either left untreated or 'multipulsed' with 10 μM BrdU or PBS. The percentage of cells in G1, S, and G2/M phases is indicated. This experiment was performed independently eight and four times for untreated and BrdU-'multipulsed' cells, respectively, with similar results; cell 'multipulsing' with PBS was performed once. **e**, Quantification of the percentage of HCT116 cells in G1, S, and G2/M phases of the cell cycle in an untreated sample or after cell 'multipulsing' with 10 μM BrdU or PBS. Median and extreme values (whiskers) are presented. n values are independent cell cultures: $n_{Control}$ = 8, $n_{Multipulse\ BrdU}$ = 4, and $n_{Multipulse\ PBS}$ = 1. S phase duration of HCT116 cells was estimated at 6.6 h by multiplying their median doubling time (12.2 h, as estimated in **b**) by their median S phase fraction (0.54). **f**, Representative western blot analysis with anti-Chk1 and anti-phospho-Chk1 (S345) antibodies of extracts from HCT116 cells either left untreated or 'multipulsed' with 10 μM BrdU. Cells treated for 24 h with 2 mM hydroxyurea (HU) were used as a positive control for Chk1 phosphorylation. Immunoblotting with anti-ß-actin was used as a loading control. This experiment was performed independently twice with similar results. **g**, Fraction of HCT116 cells recovered after 'multipulse' labelling with 10 μM BrdU relative to an untreated sample. Values are fractions of recovered cells from eleven independent 'multipulse' experiments; the median recovered fraction (black central line) is 0.52. **h**, Fraction of cells recovered after a 'multipulse' experiment with either 10 μM BrdU or PBS relative to an untreated sample.

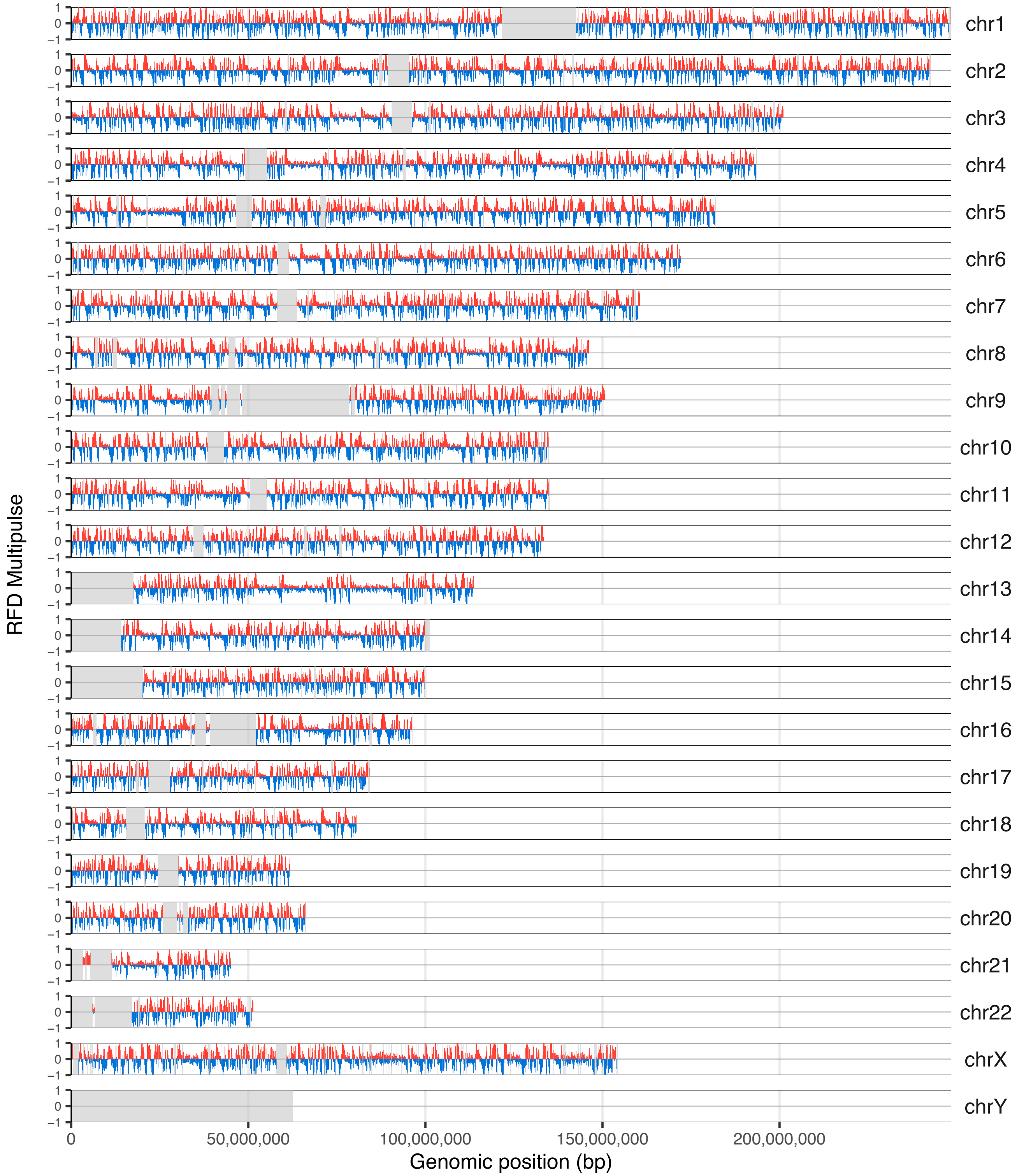


**Supplementary Figure 2. Genome-wide 'multipulse' RFD profile of HCT116 cells.** Each chromosome is presented as a separate row. Grey shaded regions mark bins excluded from downstream analyses because of abnormal read coverage (defined as low-coverage or high-coverage outlier regions, see ref. 9), and primarily correspond to centromeres. RFD is shown in 10 kb adjacent windows to allow for genome-wide plotting due to file size constraints. See Fig. 2 caption for details.
